# Metacommunity theory and metabarcoding reveal the environmental, spatial, and biotic drivers of meiofaunal communities in sandy beaches

**DOI:** 10.1101/2024.07.17.603914

**Authors:** Jan-Niklas Macher, Maximilian Pichler, Simon Creer, Alejandro Martínez, Diego Fontaneto, Willem Renema

## Abstract

Sandy beaches are important ecosystems providing coastal protection and recreation, but they face significant threats from human activities and sea level rise. They are inhabited by meiofauna, small benthic invertebrates that are highly abundant and diverse, but are commonly understudied biotic components of beach ecosystems. Here, we investigate the factors shaping meiofaunal metacommunities by employing Generalised Dissimilarity Modelling (GDM) and Joint Species Distribution Modelling (JSDM) to study community turnover and assembly processes. We analysed over 550 meiofauna samples from a >650 km stretch of the southern North Sea coastline using a metabarcoding approach. Our findings reveal that environmental factors, especially Distance from Low Tide and Sediment Grain Size, are important drivers of meiofauna community turnover. This highlights the influence of the gradient from marine to terrestrial habitats and sediment conditions. Spatial factors, which indicate dispersal limitations, also significantly impact community composition, challenging the view that marine meiofauna have broad geographic distributions. The JSDM results show that species sorting by environmental conditions is the dominant process in community assembly with increasing environmental differences between sampling sites, but that biotic associations, or similar environmental preferences, are a major driver of community assembly at sites with similar environmental conditions. Further, we find that spatial factors also significantly influence community assembly across the study region. By facilitating the inference of ecological niches for a high number of meiofaunal taxa, JSDM provides a powerful framework for understanding the ecology of these animals. Our results highlight the importance of considering environmental gradients and dispersal limitations in meiofauna and beach ecosystem research, and future research should aim at adding information on functional traits and biotic interactions under varying environmental conditions to understand meiofauna community dynamics and resilience.

## Introduction

Sandy beaches are globally common ecosystems offering crucial ecological services such as coastal protection, recreation, and supporting fisheries ^1–3^. These ecosystems face threats from “coastal squeeze”, resulting from human activities onshore and offshore, as well as sea level rise ^4–6^. At the same time, they are the least studied coastal ecosystems ^7,8^. Sandy beaches harbour diverse communities of flora and fauna ^9–11^. Among these, meiofauna, small benthic invertebrates ranging in size from about 40 µm to 1 mm ^12^, play a vital role by connecting primary producers to higher trophic levels ^13–16^. Meiofauna reach densities of a million specimens per m^2^ ^17,18^, and their known sensitivity to environmental changes also makes them potentially valuable bioindicators ^19–21^. Despite their ecological importance, a thorough understanding and study of beach meiofauna communities remains challenging due to difficulties in morphological identification and a lack of ecological knowledge ^22^. The advent of metabarcoding, a molecular technique that enables amplification and identification of many species from environmental samples, has changed meiofauna research by offering a cost-effective and efficient method of understanding meiofaunal biodiversity ^23–25^, especially when combined with morpho-taxonomic approaches^26^.

Past research has revealed a high meiofaunal species richness and notable differences in diversity within and across beaches ^9,17,27–30^. Key factors like sediment composition ^1,31^ and energy levels across the intertidal zone have been identified as important in shaping meiofaunal communities ^32–35^. Furthermore, the type and state of a beach, whether it’s reflective with a steeper slope and coarser sediment, or dissipative with a gentler slope and finer sediment, impact meiofauna community composition and diversity ^28,32,36,37^. Traditionally, marine meiofauna species were thought to have a very wide geographic distribution despite limited dispersal capabilities ^38–40^ (the “meiofauna paradox”), but this has been shown to partly stem from insufficient taxonomic resolution ^41,42^. Many meiofaunal taxa are actually limited in dispersal ^39,43^, which shapes communities and diversity on smaller and larger scales ^34,44–46^. Similarly, biological interactions such as predation and competition between meiofaunal species can significantly influence communities ^14,47,48^.

Despite these insights, the understanding of meiofaunal metacommunities remains limited. Metacommunity theory, which incorporates spatial processes and community ecology, offers a framework for studying the interplay between environmental factors, regional processes, and biotic interactions in structuring communities ^49^. Most studies on meiofauna have emphasised species sorting, which describes the process where species are filtered based on their realised ecological niches, as the most influential factor in metacommunity assembly, although dispersal limitations also play an important role ^50^. However, the application of metacommunity theory to marine and coastal meiofauna is limited. As recently highlighted in a review on major open questions in meiofauna research ^51^, there is a need for studies on spatial processes, environmental effects, and interaction in meiofauna communities.

In this study, we use Generalised Dissimilarity Modelling (GDM) to investigate how spatial and environmental factors influence community turnover in beach meiofauna (the ‘external structure’ of metacommunities ^52^), and subsequently apply Joint Species Distribution Modelling (JSDM) ^53,54^ to study the processes driving assembly of communities, and gain deeper insights into ecological niches of species (the ‘internal structure’ of metacommunities ^52^). JSDMs provide a powerful approach for inferring assembly processes shaping communities ^55–58^ and were recently enhanced to allow rapid analyses of hundreds of species and samples obtained from metabarcoding datasets ^59,60^. By providing a framework to analyse how environmental and spatial factors, as well as biotic interactions, influence communities, JSDMs can not only test existing hypotheses but also generate new, testable hypotheses about the drivers of community assembly. Here, we apply JSDMs to an extensive meiofauna metabarcoding dataset comprising over 550 samples from a stretch of >650 km of coastline of the southern North Sea.

We hypothesise that beach meiofauna metacommunities are primarily shaped by species sorting through environmental factors such as Sediment Grain Size and morphodynamic conditions, with spatial effects and species interactions playing a secondary role. By utilising GDM and the JSDM framework, our study helps understand community assembly processes and ecological niches of beach meiofauna. A better understanding of these processes and the availability of detailed information on the ecological niches of meiofauna will facilitate further research by identifying the key factors that shape biodiversity in beach ecosystems.

## Results

### Bioinformatics and OTU annotation

Following bioinformatic processing with APSCALE and quality filtering, we retained 14,822,456 mitochondrial cytochrome c oxidase I (COI) sequences and 566 Operative Taxonomic Units (OTUs). Following taxonomic annotation using NCBI GenBank and meiofauna reference barcodes reported in ^26^, we retained 11,029,442 sequences of 127 OTUs. Of these, 42 were Nematoda, 21 Copepoda, 16 Clitellata, 12 Polychaeta, 9 Gastrotricha, 8 Platyhelminthes, 7 Acoela, 4 Collembola, 3 Rotifera, 2 Branchiopoda, 1 Nemertea, 1 Tardigrada, and 1 Arachnida.

### Generalised Dissimilarity Models

The GDM explained 43.2% of the deviance, indicating the proportion of variation in community turnover accounted for by the model. The null deviance was 2223.6, and the GDM deviance was 1262.2. The intercept value was 0.9. The most influential predictor was Distance from low tide with a sum of I-spline coefficients of 1.8, with the strongest change occurring at distances corresponding to the high tide line (Figure 1 A). This was followed by Grain Size (sum of coefficients: 1.6), with the strongest change occurring between 100 and 400 μm (Figure 1 B). The third most influential factor was Geographic Distance (sum of coefficients: 0.7), with a strong change in community turnover at distances between 0 and 100km, a less pronounced change between distances of 100 to 400 km, followed by a stronger change up to 600 km (Figure 1 C). This was followed by Average Beach Slope (sum of coefficients: 0.3), with community turnover linearly increasing from flat to steep slopes (Figure 1 D), and Mean Surface Salinity (sum of coefficients: 0.2), which influenced community turnover mostly between 30 and 31 ppm (Figure 1 E). Spring Tide Range (sum of coefficients: 0.1) influenced community turnover mostly at higher ranges above 3.5 metres (Figure 1 F). Mean Annual Temperature showed a sum of coefficient of 0.0 and did not significantly influence community turnover.

**Figure 1:**
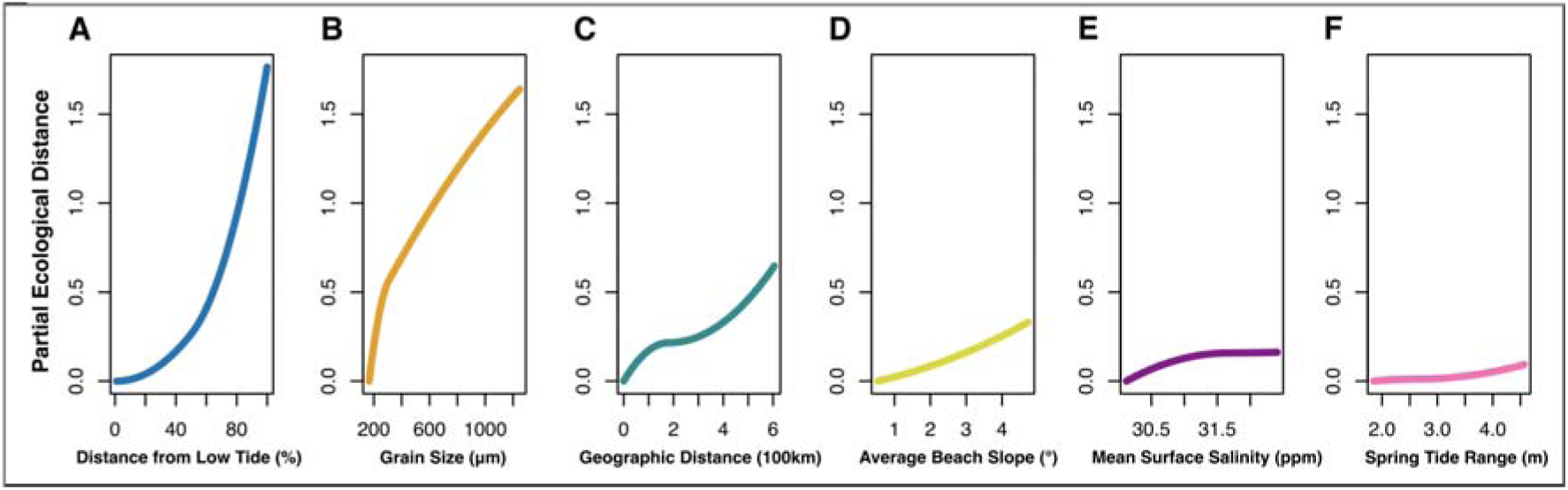
I-splines (partial ecological distance) from generalised dissimilarity modelling (GDM). The slope of the partial ecological distance indicates the rate of compositional turnover and how it changes with increasing variable values. A) Distance from Low Tide, B) Sediment Grain Size, C) Geographic Distance, D) Average Beach Slope, E) Mean Surface Salinity, F) Spring Tide Range.

### Scalable Joint Species Distribution Models

The sJSDM model resulted in a log likelihood of −5470.78 and an R^2^ value of 0.41.

### Estimated ecological niches

The distance of the sampling site from the low tide level was the most influential factor on OTU prevalence and inferred realised ecological niches. OTUs from Acoela, Branchiopoda, Copepoda, Gastrotricha, Platyhelminthes, Polychaeta, Rotifera, and Tardigrada were more prevalent closer to the low tide level, indicating a realised ecological niche in more marine conditions. In contrast, OTUs from Arachnida, Clitellata, and Collembola showed higher prevalence further away from low tide, indicating a realised ecological niche in more terrestrial conditions. Nematoda OTUs had mixed responses, with some showing higher prevalence in marine and others in more terrestrial conditions. Most OTUs exhibited a negative relationship with increasing Beach Slope, although this effect was not significant in most cases. Similarly, more OTUs showed a negative relationship with increasing Salinity. Grain size, Spring Tide Range, and Mean Annual Temperature had mixed effects on OTU prevalence, generally lacking significance (Figure 2).

**Figure 2:**
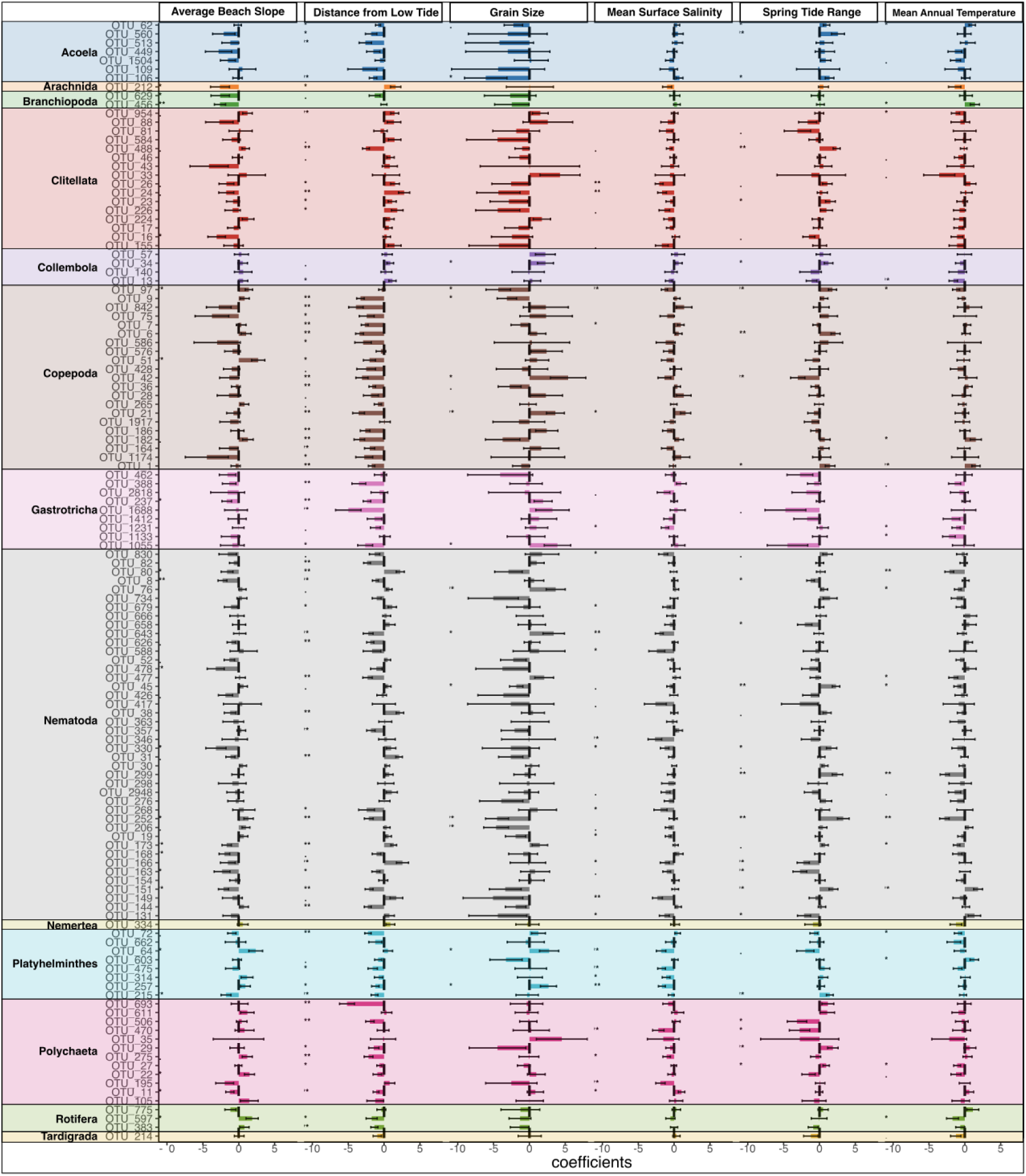
Estimated ecological niches of meiofauna OTUs. Horizontal bars show the magnitudes, directions, and standard errors of the coefficients of each of the six environmental covariates for each meiofaunal OTU, sorted by meiofaunal group. All covariates were normalised before fitting. Colours indicate meiofaunal groups. Asterisks indicate significance of the effect.

### Biotic associations

Analyses of biotic associations focusing on the top 2.5% of positive and negative correlations revealed many associations between meiofaunal OTUs. For brevity, only the most obvious results are described here. For details, please see Figure 3. Clitellata OTUs showed mostly positive correlations with other Clitellata, negative correlations with Acoela OTUs, and mixed correlations with Nematoda OTUs. Copepoda OTUs were mostly positively correlated with other Copepoda, Polychaeta, and Platyhelminthes OTUs, and showed mixed correlations with Nematoda OTUs. Nematoda OTUs showed both positive and negative correlations within their group and most other meiofauna groups, but mostly negative correlations with Gastrotricha OTUs (Figure 3).

**Figure 3:**
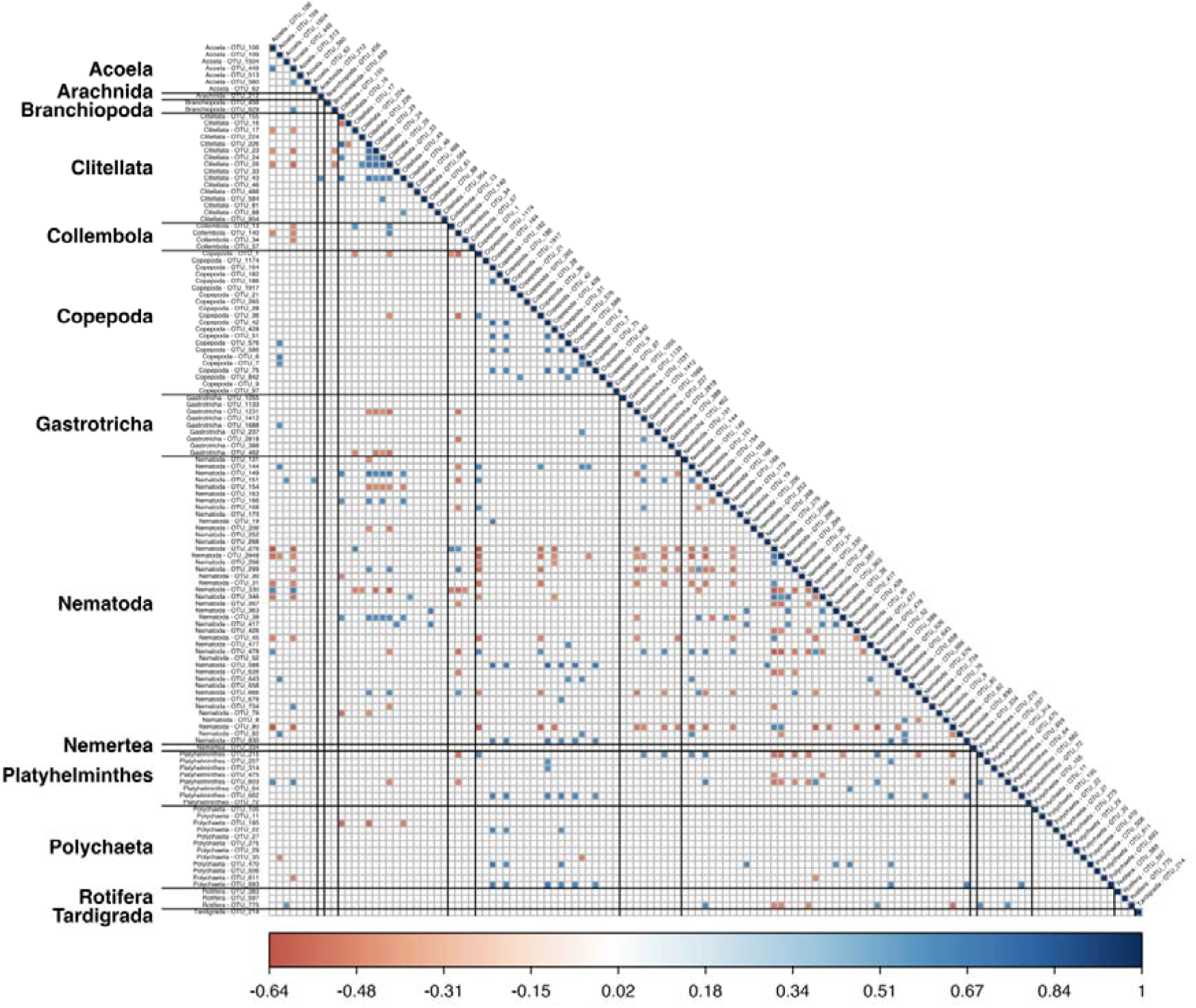
Biotic associations of OTUs by meiofaunal group, filtered to the 2.5% most negative and positive covariances. Colours indicate the magnitude of the correlation, according to the scale bar, from −0.64 to +1.00.

### The role of environmental distinctiveness for community assembly

The JSDM analyses of drivers of community assembly showed that as the environmental distinctiveness of sampling sites increased, the R² explained by the environmental component rose significantly and linearly from low to high environmental distinctiveness, indicating stronger species sorting due to environmental differences in sampling sites. The biotic covariance component showed a strong non-linear response with increasing environmental distinctiveness, dropping from low to medium levels, before increasing towards high levels of environmental distinctiveness. This indicates that biotic interactions or similar environmental preferences of meiofaunal OTUs are more important both at low and at high levels of environmental distinctiveness of sampling sites (Figure 4 A).

**Figure 4:**
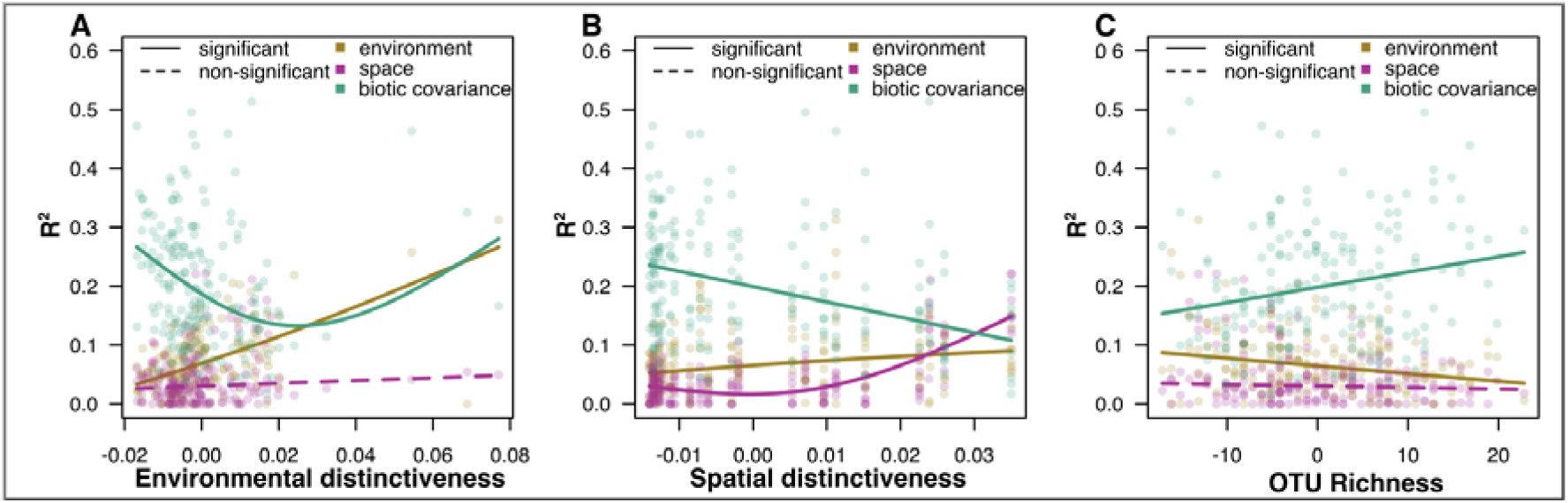
Correlation of the importance of assembly processes to environmental predictors. Quantile regressions correlating the importance of the three assembly mechanisms environment (yellow), space (purple), and biotic covariance (green), measured by the share of absolute partial R^2^ values, per sampling site against A) environmental distinctiveness, B) spatial distinctiveness, C) OTU Richness. Significant effects (p <0.05) are shown as continuous lines, non-significant effects are shown as dashed lines.

### The role of spatial distinctiveness for community assembly

With increasing spatial distinctiveness of sampling sites, the R² explained by the spatial component increased significantly from medium to high levels, indicating stronger dispersal barriers between sampling sites that are further apart. At the same time, the environmental component also increased slightly with increasing spatial distinctiveness. The biotic covariance component decreased significantly from low to high levels of spatial distinctiveness (Figure 4 B).

### The role of OTU richness for community assembly

With increasing OTU richness per sampling site, the R² explained by biotic covariance showed an increase from low to high OTU richness, indicating stronger biotic associations of OTUs. At the same time, the environmental component decreased significantly with increasing OTU richness (Figure 4 C).

### Influence of individual environmental covariates on community assembly

The R^2^ for the environmental component increased significantly (p <0.05) with increasing Grain Size, indicating that coarser sediments play an important role in meiofauna community assembly (Figure 5A). With increasing Distance from Low Tide, only the biotic association component showed a strong nonlinear relationship, peaking at intermediate distances from low tide (Figure 5B). With increasing Spring Tide Range, we found a significant, but minor increase in the R² value of the environmental component, particularly between low and medium values. The spatial component also increased, but explained less than the environmental component. The biotic association component declined significantly from low to high Spring Tide Range values (Figure 5C). With increasing Beach Slope, the environmental component increased significantly from intermediate to high values. The spatial component showed a slight increase (Figure 5D). Mean Surface Salinity did not significantly change the environmental, spatial or biotic covariance component (Figure 5E). For the Mean Annual Temperature, we found a decrease in the spatial component from low to intermediate levels of Annual Temperature. The biotic association component showed a non-linear relationship with increasing temperature, with lowest values at low and high temperatures and a peak at intermediate levels (Figure 5F).

**Figure 5:**
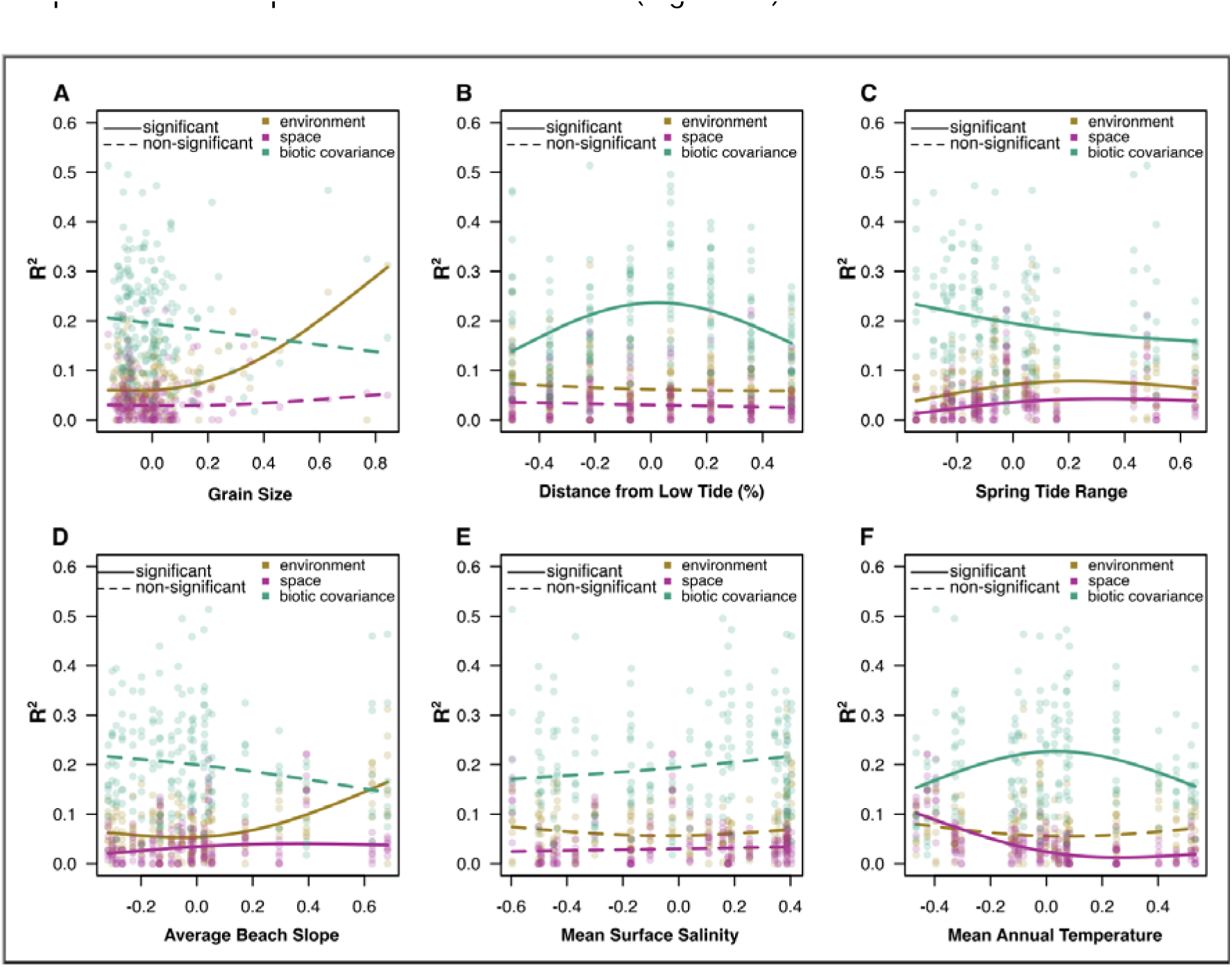
Correlation of the importance of assembly processes to the six environmental covariates. Quantile regressions correlating the importance of the three assembly mechanisms environment (yellow), space (purple), and biotic covariance (green), measured by the share of absolute partial R² values, per sampling site against (A) Grain Size, (B) Distance from Low Tide, (C) Spring Tide Range, (D) Average Beach Slope, (E) Mean Surface Salinity, (F) Mean Annual Temperature. Significant effects (p < 0.05) are shown as continuous lines, non-significant effects are shown as dashed lines.

## Discussion

We studied the processes shaping sandy beach meiofauna metacommunities and hypothesised that species sorting is primarily driven by environmental conditions due to the strong gradient from marine to terrestrial habitats on beaches, and to a lesser extent by spatial factors and biotic interactions. We applied Generalised Dissimilarity Modeling (GDM) to study the external structure of the beach meiofauna community, i.e, the community turnover as a result of environmental and spatial factors, and Joint Species Distribution Modeling (JSDM) to study the internal structure of the metacommunity, i.e., the environmental, spatial, and biotic drivers of community assembly. Our results support that species sorting is a key process in beach meiofauna community assembly, as both GDM and JSDM analyses revealed that environmental factors are major drivers of community turnover and assembly. This finding aligns with the species sorting aspect of metacommunity theory and confirms common findings that meiofauna are strongly influenced by the physical factors of their habitat ^1,28,32,33,61^.

### Community turnover

The GDM results show that, in line with our expectations, the environment is the major driver of community turnover in sandy beaches, with the Distance from Low Tide line being the main factor. The strongest change in community turnover is found at a Distance from Low Tide that corresponds to the high tide level, indicating that the transition from mostly marine to mostly terrestrial conditions, which includes major changes in salinity, humidity, and temperature, is the main driver of community turnover. This corresponds to previous findings on beach invertebrate communities, which found a clear differentiation of communities from supralittoral and intertidal areas of the beach ^62^. The second most important environmental factor was Sediment Grain Size, which is in line with previous studies that found major impacts of grain size on meiofauna communities ^28,32,63^, as grain size affects habitat structure by influencing, among others, interstitial space availability and sediment oxygenation.

The spatial factor, i.e., the distance between sampling sites, was the third most important variable, showing that dispersal limitations play a role in beach meiofauna in the study region. Previous studies on dispersal of meiofauna in marine systems found strong differences between taxonomic groups ^64,65^, with higher local endemicity of e.g. Copepoda ^66^, and less pronounced biogeographic structure for Nematoda ^67^. To our knowledge, while the dispersal of intertidal meiofauna has been studied (e.g.^68,69^), there is no such study for supralittoral meiofauna, which would be important to understand beach meiofauna communities holistically. The fourth important factor for community turnover was Beach Slope, which reflects the morphodynamic conditions on a beach and, a factor known to be important in shaping communities ^70^. In our study area, however, it seems that this factor is less important, or its effects are overshadowed by other environmental factors, potentially because most beaches in the study area are of an intermediate or dissipative type. These differences might be more pronounced between reflective and dissipative beaches.

The remaining environmental factors, Mean Surface Salinity and Spring Tide Range, played minor roles in driving community turnover. The strongest change in community turnover attributed to salinity was found between low and medium salinity levels, which is in line with findings showing that many meiofauna species are sensitive to changes in salinity ^71,72^. Spring Tide Range, in contrast, influenced community turnover only at high Spring Tide Range, indicating that the higher energy levels connected to higher Spring Tide Ranges are not very influential in the study area, despite generally being regarded as an important factor in beach ecology ^35^. Mean Annual Temperature did not influence community turnover in the study area, potentially indicating that there are relatively homogeneous meteorological conditions across the study area. However, temperature differences might play a role on a smaller scale, e.g. due to different currents, beach geomorphology, influx of coastal groundwater, or different sunlight reflection on beaches depending on grain size, seasonality and exposure times ^73^.

### Community assembly processes

The JSDM results revealed more complex patterns. Our results show that environmental factors strongly influence community assembly, especially in sites with distinct environmental conditions. This finding supports our hypothesis that species sorting is an important driver of meiofauna community assembly, confirming the assumptions of Leibold et al. ^52^. However, we also observe that in samples from similar environmental conditions, biotic interactions between meiofauna or similar environmental preferences play a more important role, while increasing environmental differences filter species according to their realised ecological niches. This underscores the importance of considering both biotic and abiotic factors when studying beach meiofauna communities and their response to the environmental conditions.

Of the individual environmental variables, grain size emerged as the most influential factor driving community assembly, consistent with the GDM results showing its strong impact on community turnover. The optimum grain size for intertidal meiofauna in our study area, where the environment is not a limiting factor for most meiofauna, appears to be at the lower end of the spectrum. Since the median grain size in our dataset was 300 µm, this supports the assumption that an optimum grain size for interstitial fauna, with an ideal balance of interstitial space, water retention, oxygenation, and organic content retention, lies between 200 and 400 µm^70^.

Unexpectedly, the importance of the environmental component in community assembly did not increase with distance from the low tide line, despite being a primary driver of community turnover in the GDM analyses, and also emerging as a major factor in the analysis of realised ecological niche per OTU. Instead, the biotic association component was most influential at intermediate distances from low tide, corresponding to the area of the high tide line. This suggests that strong community turnover might result from the exclusion of taxa at the marine-terrestrial intersection and the co-occurrence of meiofaunal species in these distinct habitats, that might overshadow any environmental effects. A more detailed study of within-beach diversity patterns, including a higher number of environmental variables measured across the intertidal zone, such as pH, salinity, and organic content of the sediment, might allow deeper insights of the drivers of the observed pattern.

We found that steeper slopes were more influential in defining community assembly, which indicates that energy levels and habitat stability differ significantly between steep and less steep beaches, thereby leading to increased species sorting. Beach Slope influences community assembly due to strong morphodynamic influences, with steeper slopes serving as stronger environmental filters for meiofauna ^1,28,70,74^, and the optimum for meiofauna is thought to be found in intermediate conditions ^70^. We found that for Spring Tide Range, the strongest change in explanatory power of the environmental component was found between low and medium ranges, suggesting that the change in hydrodynamic forces between and medium ranges is most important for filtering meiofauna.

Salinity did not significantly influence community assembly, even though the GDM analyses indicated a minor influence on community turnover, potentially indicating that other environmental factors overshadow the effect of salinity on community assembly. Other studies found a strong influence of salinity on meiofauna ^72,75,76^, and we suggest that smaller scale changes in salinity across the intertidal zone might be more influential in shaping communities ^77,78^. Mean Annual Temperature mainly affected the biotic associations, indicating more species co-occur at intermediate temperatures, which we interpret as reflecting similar realised ecological niches and thereby higher co-occurrence. That the spatial component was more important at low Annual Temperature seems plausible, as the lowest temperatures are found in the northern range of our study area, which are also furthest away from the other sampling sites.

### Spatial factors and biotic association

While species sorting was the dominant factor, spatial factors also played a significant role in community assembly and indicated dispersal limitations with greater distances between sampling sites. The observed spatial structure suggests that while meiofauna can disperse across beaches, the dispersal is insufficient to entirely homogenise communities over the approximately 650 km of coastline included in our study. This finding further challenges the traditional view that meiofauna have broad geographic distributions and underscores the importance of considering dispersal limitations in future meiofauna studies ^39,79^. However, our results also indicated an interaction between spatial and environmental factors, with spatially unique sites being environmentally more distinct. This is plausible, as some sampling sites towards the northern limit of our study area were also among the sites with the coarsest grain size and steepest Beach Slope, while the southernmost sampling sites were among the ones with the shallowest Beach Slope. Since this is inevitable in natural ecosystems, we highlight the need for running controlled experiments to understand the interplay between local and regional processes and environment, e.g., by running controlled field experiments ^80,81^ in different geographic areas to gain a better understanding of the interaction of spatial and environmental factors.

Our JSDM results highlighted that biotic associations play an important role in community assembly, especially in environmentally similar sites. We found that taxa known to occur in the same habitat exhibit the strongest co-occurrence, such as Copepoda, Platyhelminthes, and Polychaeta, which are primarily marine taxa. The decrease in biotic associations with increasing environmental distinctiveness supports the idea that species interactions in beach ecosystems are more pronounced in homogeneous environments ^82^, where niche differentiation and competitive exclusion can occur. However, we note that without sufficient information on the ecological traits of most meiofaunal species, the nature of these interactions remains speculative. Moreover, biotic associations in JSDM can also arise from missing environmental predictors ^83^.

### Implications for Future Research

Our study provides deeper insights into the realised ecological niches of sandy beach meiofauna and the factors shaping their communities. Future research should aim to investigate a broad range of meiofaunal taxa to understand their traits, as a combination of morphological, physiological, and behavioural traits will help link traits with the ecological roles of species. Previous studies on selected taxa have highlighted the importance of biotic interactions among meiofauna species ^14,47,48^. Metabarcoding and JSDM analyses facilitate the study of meiofauna niches through high-throughput sequencing, but more studies on selected species are necessary to verify results and understand interactions. Specifically, research should focus on how factors such as body size, feeding type, reproductive strategy, and the interaction of species and traits influence meiofauna distribution along the marine-to-terrestrial gradient and across large geographic scales.

Controlled field and mesocosm experiments are needed to isolate specific variables and test the relationships between environmental and spatial factors in community assembly. Although replicating natural beach conditions while manipulating variables like sediment type, salinity, and slope is challenging due to the dynamic nature of littoral ecosystems, such experiments can validate findings and enhance understanding of meiofauna responses to different environmental conditions ^84–87^. For example, experiments with selected pairs of species or communities could test the hypothesis that species interactions are more significant near the high tide line than in other zones due to unique environmental conditions, but could also test whether interactions are more important under harsher environmental conditions and increasing stress, as has been suggested for sandy beach macrofauna ^88^.

Moreover, more studies on dispersal limitations in various meiofaunal groups are needed to understand dispersal mechanisms and colonisation dynamics. Identifying the factors that limit dispersal is crucial for informing conservation strategies. Hypotheses regarding the significance of geographic and environmental barriers in limiting meiofauna dispersal could be tested by tracking dispersal events of specific species over varying geographic scales ^79,89^. For example, providing specific habitat conditions at different intervals from source populations ^46^ and using connected micro-or mesocosms separated by areas with potentially adverse or facilitating environmental conditions could test these ideas. Finally, molecular methods enable the rapid analysis of hundreds of samples. While they need rigorous testing and verification ^90–92^, they are powerful tools in combination with statistical methods such as JSDM, facilitating analyses of larger and more complex datasets. For beach ecosystems and meiofauna communities, this enables more comprehensive studies of biodiversity across the entire beach ecosystem, and could help determine the often hidden biodiversity of beach animals and their ecology, but can also help address ecological questions such as to which extent beaches can be considered closed or semi-closed ecosystems, as suggested by McLachlan et al. ^93^.

## Conclusion

Our study supports the hypothesis that beach meiofauna metacommunities are primarily shaped by environmental conditions, with spatial effects and biotic associations also playing important roles. The findings deepen the ecological understanding of processes shaping these metacommunities and highlight the need for further research on species interactions and functional traits driving assembly processes. By integrating high-throughput sequencing data with statistical modeling, JSDM analyses offer a framework to understand complex relationships between environmental variables, spatial factors, and biotic interactions, providing insights into biodiversity in sandy beach ecosystems and beyond.

## Material & Methods

### Sampling and environmental variable measurement

We collected meiofauna from 24 sea-facing, unsheltered sandy beaches of the southern North Sea, covering 650 km of coastline between Zeeland (southern Netherlands) and Sylt (northern Germany). Sampling took place during the summers of 2021 and 2022 (See Supplementary Table 1 for coordinates and Fig. 6A for a map of sampling sites).

**Figure 6.**
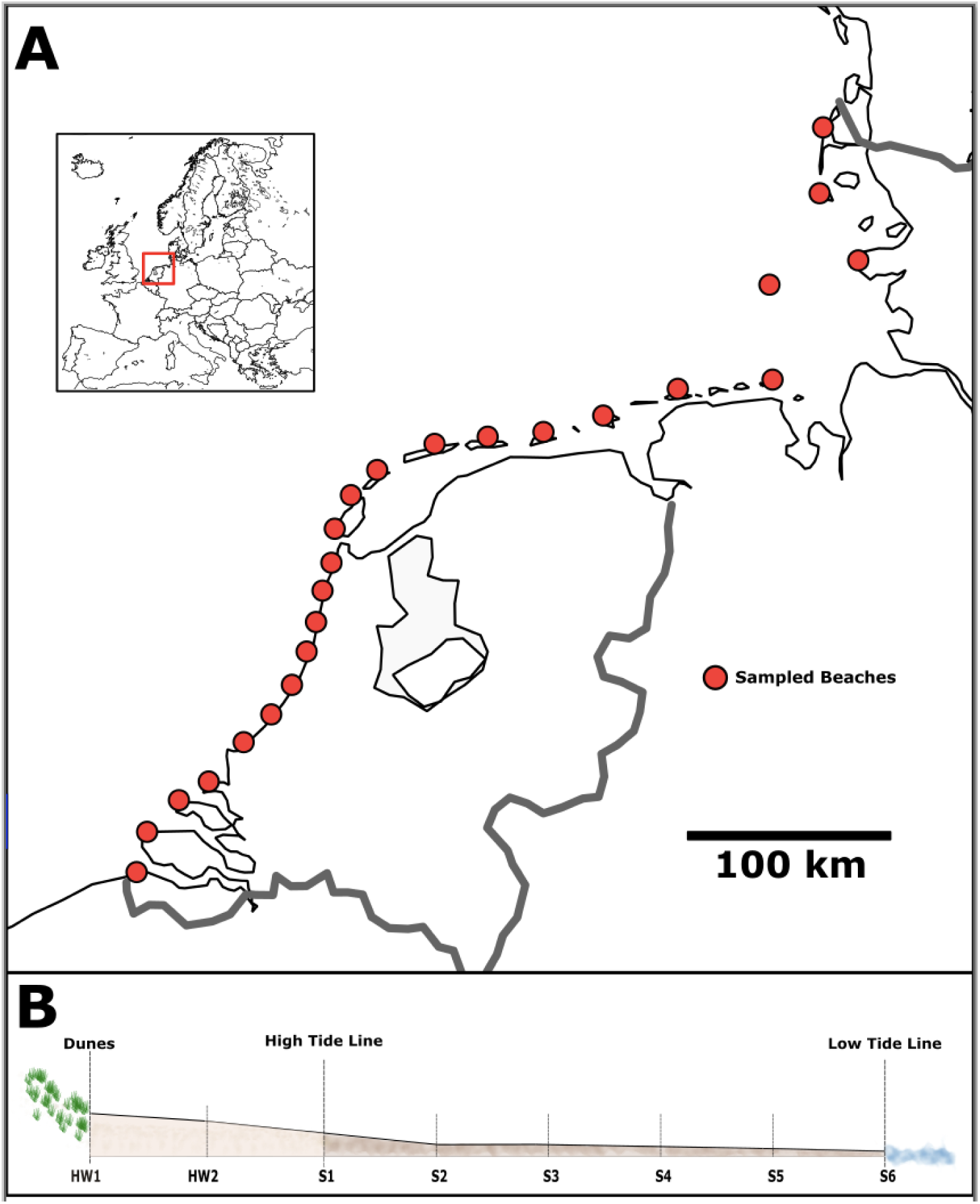
A) Map of the study area showing the 24 sampled beaches. The small map indicates the location of the study area in Europe. B) Schematic view of a sampling transect across a beach.

Samples were taken during daytime and at maximum low tide. We sampled along three parallel transects per beach, each with eight sampling sites. The first sample was taken at the foot of the dunes, the second sample halfway between the dunes and the high-tide line, and six samples were equidistantly spaced from the high-tide line to the low-tide line (see Fig. 6B). Sampling followed established protocols for the intertidal zone of sandy beaches, including the measurement of beach width from high tide line to low tide line, measuring of the beach slope (in degrees), wave period (in seconds) and breaker height (in metres) ^35^. The tidal range was extracted from online databases (www.tide-forecast.com), and we calculated the Relative Tide Range (RTR) index for each beach ^35^. Furthermore, we assessed the beach state (reflective, intermediate, dissipative) ^1^. At each sampling site, we collected two sediment cores using sterile plastic syringes: One core of 5 cm diameter and a length of 10 cm (volume ≈200 ml), and a second core of 1 cm diameter and a length of 10 cm (volume ≈8 ml). The small sediment core was immediately transferred to a 50 ml Falcon tube and the large sediment core was transferred to a sterile 1l plastic bottle. We extracted meiofauna from the large sediment core (≈200 ml) directly on the beach, using the MgCl2 decantation method ^94^. We added 500 ml of isosmotic MgCl_2_ solution to the sediment, which anaesthetizes meiofauna and allows their separation from the sediment by decantation. After 5 minutes, the sediment in the MgCl_2_ solution was carefully swirled ten times, and the supernatant containing meiofauna was decanted through a 1 mm and 41 µm sieve cascade, as commonly done in beach meiofauna studies ^95–97^. The meiofauna fraction retained on the 41 µm sieve was rinsed into sterile 15 ml Falcon tubes and preserved with 10 ml 96% EtOH. All sampling equipment was thoroughly rinsed with ethanol after taking each sample to prevent contamination. All samples were transported back to the Naturalis Biodiversity Centre laboratory and stored at −20 °C until further processing. Sediment from the smaller core was dried and the grain size was measured on a LS13320 Particle Size Analyzer (Beckman-Coulter, USA) for the eight samples of the central transect per beach.

### DNA extraction, amplification and sequencing

We extracted DNA from dried meiofauna samples after evaporating the ethanol at 50 °C overnight in a sterile warming cabinet and transferring the dried samples to 2 ml Eppendorf tubes. DNA extraction was performed using the Macherey Nagel NucleoSpin Soil kit (Macherey Nagel, Düren, Germany) following the standard protocol including bead beating, but with an additional overnight Proteinase K digestion step (50 µl 250 µg/ml ProtK, Thermo Fisher Scientific, Waltham, USA) added to the lysis buffer provided with the kit to improve cell lysis, as done in previous studies on meiofauna ^95,98^.

For community metabarcoding, we amplified meiofauna DNA using a two-step PCR protocol with the widely-used LerayXT primers targeting a 313 base pair region of the mitochondrial cytochrome c oxidase I (COI) gene of a broad range of Eukaryota ^99,100^. The first PCR reaction contained 11.7 µl mQ water, 2 µl Qiagen CL buffer (10x; Qiagen, Hilden, Germany), 0.4 µl MgCl2 (25 mM; Qiagen), 0.8 µl Bovine Serum Albumin (BSA, 10 mg/ml), 0.4 µl dNTPs (2.5 mM), 0.2 µl Qiagen Taq (5U/µl), 1 µl of each nextera-tailed primer (10 pMol/µl), and 2.5 µl of DNA template. PCR amplification involved an initial denaturation at 96 °C for 3 minutes, followed by 30 cycles of denaturation for 15 seconds at 96 °C, annealing at 50 °C for 30 seconds, and extension for 40 seconds at 72 °C, concluding with a final extension at 72 °C for 5 minutes. We processed six negative controls (containing Milli-Q water instead of DNA template; Milli-Q, Merck, Kenilworth, USA) alongside the samples to check for potential contamination.

After the first PCR, samples were cleaned with AMPure beads (Beckman Coulter, Brea, United States) at a 0.9:1 ratio according to the protocol to remove short fragments and primer dimers. The second PCR involved amplification with individually tagged primers, following the same protocol as above and using the PCR product from the first PCR as the template, but reducing the PCR cycle number to 10. We measured DNA concentrations using the FragmentAnalyzer (Agilent Technologies, Santa Clara, CA, USA) with the High Sensitivity Kit and pooled samples equimolarly. The final library was cleaned with AMPure beads as described above and sent for sequencing on three Illumina MiSeq runs (2 × 300 bp read length) at Baseclear (Leiden, The Netherlands).

### Bioinformatic processing of community metabarcoding data

We processed the raw metabarcoding reads using APSCALE ^101^ with the following settings: maximum differences in percentage: 20; minimum overlap: 50, minimum sequence length: 310 bp; maximum read length: 316 bp, minimum size to pool: 20 sequences. Sequences were clustered into Operational Taxonomic Units (OTUs) with a sequence similarity threshold of 97%. To account for potential low-level contamination or tag jumping common on Illumina platforms ^102^, we removed OTUs with an abundance of <0.03% of reads per sample, following a common subsetting approach in metabarcoding studies ^103,104^. We removed 11 samples with less than 3000 reads per sample during bioinformatic processing, retaining samples for which 1 read corresponds to >0.03% of the total reads number. We performed taxonomic assignment using NCBI GenBank expanded with meiofauna COI barcodes generated as part of associated taxonomic workshops in Leiden ^26^. Taxonomic ranks were assigned to OTUs using established identity thresholds: >97%: species, >95%: genus, >90%: family, >85% order ^105^. Taxonomic annotation of the reads present in the six negative controls (5,211 reads total) showed that only two OTUs (OTU222, *Navicula,* diatom, with 2,728 reads; and OTU219, *Homo sapiens*, with 2,653 reads) were dominant in negative controls, and these OTUs were subsequently excluded from the dataset. Further, following ^106^, a strict read filtering was applied, by subtracting the sum of reads per OTU present in negative controls from the reads per OTU in each sample. Merging the three biological replicates per tidal level and beach resulted in 190 composite samples.

Subsequently, OTUs that were assigned with less than 85% identity to a reference or identified as non-meiofauna taxa were excluded from further analyses. We merged the three replicates per tidal level per beach into one composite sample to account for potential variability within tidal levels, resulting in 190 composite samples. Following this, we further retained only OTUs that were present in at least 10 samples, following the recommendation of Cai et al. ^107^, since sjSDM can be sensitive to false negative occurrences.

### Environmental variable selection and Generalised Dissimilarity Modelling

We measured the following environmental variables: Grain Size (average per sample, in μm), Distance from Low Tide Level (in percentage, with low tide = 0 and the sample closest to the dunes = 100 percent), Beach Width (in metres), Salinity of the Surface Water (in ppm), Spring Tide Range (in metres), Average Beach Slope (in degrees). Further, we obtained the following variables from Bio-ORACLE v2 database containing marine data for modelling ^108^: Mean Surface Water Temperature (°C), Mean Surface Phosphate (mmol m^−3^), Mean Surface Nitrate (mmol m^−3^), Surface Dissolved Oxygen (mmol m^−3^), Mean Surface Salinity (PSS). We also downloaded bioclimatic variables from the WorldClim 2 database ^109^, which contains data for terrestrial environments: Annual Mean Temperature (°C) and Annual Precipitation (mm). All variables were extracted from the data layers using QGIS v. 3.36 (http://qgis.org/). We calculated pairwise correlation of the variables using the cor function in R and removed variables that showed a correlation coefficient > 0.7. Following this, the following variables were left: Grain size, Distance from Low Tide, Beach width, Spring Tide Range, Average Beach Slope, Mean Primary Production, Mean Surface Salinity, Average Annual Temperature. We then used the collinearity diagnostic variance inflation factor, implemented in the ‘vifstep’ function of the R package usdm ^110^ with a threshold of 0.5, which excluded the variables Mean Primary Production and Beach Width. The retained variables, which all showed a VIF score < 2, were Grain Size, Distance from Low Tide, Spring Tide Range, Average Beach Slope, Mean Surface Salinity, and Average Annual Temperature. We used the R package ‘gdm’ ^111^ to calculate the GDM based on a Jaccard distance matrix of community similarity. The environmental variables were provided as predictors, and geography, based on a distance matrix, was included in the calculations. The GDM plots were generated with the plot function from the gdm package.

### Scalable joint species distribution models (sjSDM)

We fitted the sJSDM model using the ‘sjsdm’ package v.1.0.5 ^53^ in R. We modelled the spatial relationships between sites as a polynomial of second order of the scaled coordinates (trend surface model) ^112,113^. All OTUs were converted to presence/absence and fit with a binomial model with probit link. We modelled the responses to environmental gradients as linear responses, set the learning rate scheduler to 10, the reduce factor to 0.9 (optimizer parameters that help with the convergence), and ran the model with 500 iterations. We visualised the estimated effects of the six environmental covariables on OTU prevalence using the model plot function in the sjsdm package, with OTUs coloured and sorted by meiofaunal groups. Further, we plotted the biotic associations of OTUs (the estimated variance-covariance matrix, normalised to a correlation matrix), sorted by meiofaunal groups, by extracting the correlation coefficients per pair of OTUs from the sjSDM model output and calculating pairwise correlation plots after filtering to the 2.5% most negative and positive values. We used variation partitioning to calculate the importance of the three assembly processes “environment”, “space”, and “biotic association” per sampling site and per OTU, the internal structure^52^, and we regressed the individual sample R^2^ values for “environment”, “space”, and “biotic covariance” against the environmental distinctiveness of samples using quantile regression (50% quantile) because of the non-normal distribution of the R^2^ values and potential outliers. We also tested the same processes against spatial distinctiveness of samples, and also against OTU richness per sample. Furthermore, we tested each environmental covariate individually.

